# Personalized Perturbation Profiles Reveal Concordance between Autism Blood Transcriptome Datasets

**DOI:** 10.1101/2021.01.25.427953

**Authors:** Jason Laird, Alexandra Maertens

## Abstract

The complex heterogeneity of Autism Spectrum Disorder (ASD) has made quantifying disease specific molecular changes a challenge. Blood based transcriptomic assays have been performed to isolate these molecular changes and provide biomarkers to aid in ASD diagnoses, etiological understanding, and potential treatment^1–6^. However, establishing concordance amongst these studies is made difficult in part by the variation in methods used to call putative biomarkers. Here we use personal perturbation profiles to establish concordance amongst these datasets and reveal a pool of 1,189 commonly perturbed genes and new insights into poorly characterized genes that are perturbed in ASD subjects. We find the resultant perturbed gene pools to include the following unnamed genes: C18orf25, C15orf39, C1orf109, C1orf43, C19orf12, C6orf106, C3orf58, C19orf53, C17orf80, C4orf33, C21orf2, C10orf2, C1orf162, C10orf25 and C10orf90. Investigation into these genes using differential correlation analysis and the text mining tool Chilibot reveal interesting connections to DNA damage, ubiquitination, R-loops, autophagy, and mitochondrial damage. Our results support evidence that these cellular events are relevant to ASD molecular mechanisms. The personalized perturbation profile analysis scheme, as described in this work, offers a promising way to establish concordance between seemingly discordant expression datasets and expose the relevance of new genes in disease.

## Introduction

ASD is an incredibly complex disease marked by an even more complex genetic presentation. The Simons Foundation Autism Research Initiative SFARI has curated a list of over 900 genes implicated in ASD^7^. Males seem more likely to develop ASD, with Loomes et al. 2017 noting the ASD male to female ratio is close to 3:1^8^. This discrepancy between males and females is not currently understood^9^. What is more, there is an extreme bias towards males in autism research with Ratto et al. 2019 noting that female specific ASD traits are being missed by behavioral diagnosis^10^. Multiple transcriptomic studies have been performed with the aim of supplementing psychiatric data with quantification of the molecular differences between typically developing individuals and individuals with ASD^1,4–6,11,12^. The transcriptomic assays examined in this work are blood-based, which offers the advantage of being easily available but the disadvantage of only revealing indirectly the likely phenotypic changes seen in the brain. Brain biopsies are more difficult to obtain, and are typically a complex mix of cells, whereas blood is more readily available and can be tracked over time. By exploring aberrant genetic expression in blood, we aim to offer a more robust understanding of ASD molecular mechanics in a tissue well suited for sampling.

Instead of traditional methods of establishing differential expression, this work uses “personal perturbation profiles” – wherein a z-score based on the expression values of the control group determines whether a gene in an ASD subject is perturbed^13^. Personal perturbation profiles are an admittedly interesting choice for a concordance study as Menche et al. 2017 showed these profiles often have very little overlap, even in the same study^13^. However, traditional methods of establishing differential expression are not without drawbacks. Isolating a common pool of biomarkers across studies is often difficult due to the variety in platforms and statistics used in delivering these results^13^. Balázsi et al. 2011 point out biological processes are marked by random fluctuations providing an inherent source of noise when analyzing biological data^14^. While these concerns are not explicitly resolved with personal perturbation profiles, they do attempt to account for another source of biomarker variation between studies, disease heterogeneity and a disease that results from a multitude of molecular mechanisms^13^. Given the apparent heterogeneity of ASD, the personal perturbation approach seemed appropriate.

This work aims to reveal not only a set of commonly perturbed genes implicated in ASD but explore poorly characterized genes that are commonly perturbed in ASD individuals. We selected genes with no canonical name at the time these studies were performed. These genes were isolated by the pattern “CXXorfXXX” in the gene name. While isolating genes based on this pattern is a relatively quick and simple way of isolating poorly characterized genes, it has some advantages over other methods – for example, we could have selected genes based on PMID count or GO annotations. However, the majority of GO annotations are held by only 16% of all known genes,^15^ and these annotations are used as evidence in many articles which may inflate literature counts. Additionally, not all genes are mapped to PMIDs, nor is there any obvious cut-off as to what makes a gene poorly studied. For the sake of a first pass analysis, we chose to pick unnamed genes to isolate a smaller pool of poorly characterized candidates.

## Results

### Personalized Perturbation Profiles Reveal a Set of Commonly Perturbed Genes

The following ASD blood-based transcriptomic studies were examined, GSE26415,GSE6575,GSE37772, GSE18123-GPL570,GSE18123-GPL6244,GSE42133, and GSE25507. Data was pulled from the Gene Expression Omnibus GEO. The tissue source used in GSE26415, GSE6575, GSE37772, GSE18123-GPL570,GSE18123-GPL6244, GSE42133, and GSE25507 were venous leukocytes, whole blood, lymphoblast cell line, blood, blood, leukocyte, and lymphocytes, respectively. Previous studies into ASD blood-based transcriptomics have shown via multidimensional scaling that there are notable differences between these tissue types^16^. However, the availability of large blood based ASD datasets has limited us to these datasets in this first attempt to establish concordance.

We first began by taking a more traditional approach to differential expression, by using the Welch two sample t-test to isolate which genes were differentially expressed in each dataset. T-test p-values were limited to those below the canonical 0.05 threshold to reveal each dataset had an average of 4,436 differentially expressed genes. When each dataset’s differentially expressed genes were compared with one another, these datasets only had one gene in common, SMARCA2.

To establish personal perturbation profiles, the expression of each gene for each ASD individual was compared to the expression values of the controls for that gene per study. If the z-score of the expression value for the ASD individual was greater than 2.5 or less than −2.5 the gene was considered perturbed. The list of perturbed genes per patient was added to the list of genes perturbed in that study (**Supplemental Data 1**). On average there were 14,117 unique personally perturbed genes per study **Figure 1**. These pools were compared with one another to reveal 1189 genes in common **Figure 1** (**Supplemental Data 2**). In total, there were 534 ASD individuals between the studies examined. Only 9.2% of these ASD individuals were female and the remaining 90.8% were male. The extreme bias toward males in these data prompted us to account for gender. To obtain a pool of male specific personally perturbed genes each study’s list of personally perturbed genes was limited to the genes found to be perturbed in male ASD patients. These male only perturbed gene pools were compared with one another to obtain a pool of commonly perturbed male genes. The pool of commonly perturbed female genes was obtained using this same scheme. ASD males were found to have 1,111 commonly perturbed genes (**Supplemental Data 2**), while ASD females were found to have 133 commonly perturbed genes (**Supplemental Data 2**).

**Figure 1.**
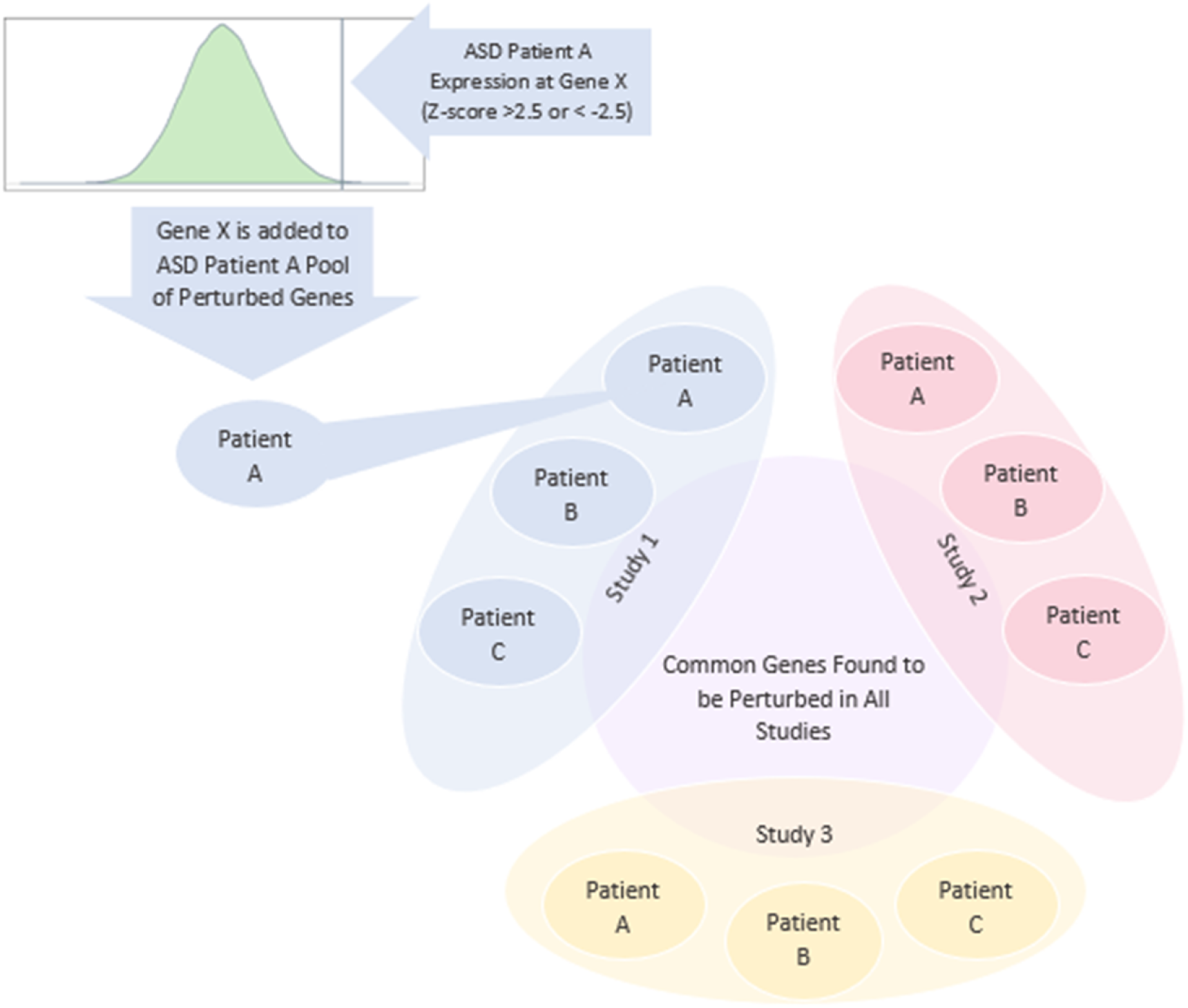
Pools of personally perturbed genes per study reveal a set of commonly perturbed genes. The expression of each gene for each ASD individual was compared to the expression values of the controls for that gene per study. If the z-score for the expression of the ASD individual was greater than 2.5 or less than −2.5 the gene was considered perturbed. The list of perturbed genes per patient was added to the list of genes perturbed in that study. These pools of genes were compared with one another to reveal a set of 1189 commonly perturbed genes, 1,111 genes when each study’s pool was limited to males and 133 when the pool was limited to females.

### PANTHER Enrichment of Personally Perturbed Genes

The common personally perturbed genes, male specific personally perturbed genes and female personally perturbed genes were run through the PANTHER Overrepresentation Test GO Biological Process Complete, GO Molecular Function Complete, and GO Cellular Component Complete utilizing the GO Ontology database (DOI: 10.5281/zenodo.4081749 Released 2020-10-09) (**Supplemental Data 3, Supplemental Data 4, Supplemental Data 5, Supplemental Data 6, Supplemental Data 7, Supplemental Data 8, Supplemental Data 9, Supplemental Data 10, Supplemental Data 11**)^17^. Out of the set of 1189 personally perturbed genes with no gender filter, 21 of these genes did not map to any annotations. When the male and female specific genes were examined, 18 and 2 genes did not map to any annotation, respectively. These annotations were filtered to exclude any annotation with more than 1000 genes linked to it. In this way we avoid overly ambiguous annotations. The top 10 most significant annotations for each category per pool of genes is displayed in Figure 2.

**Figure 2.**
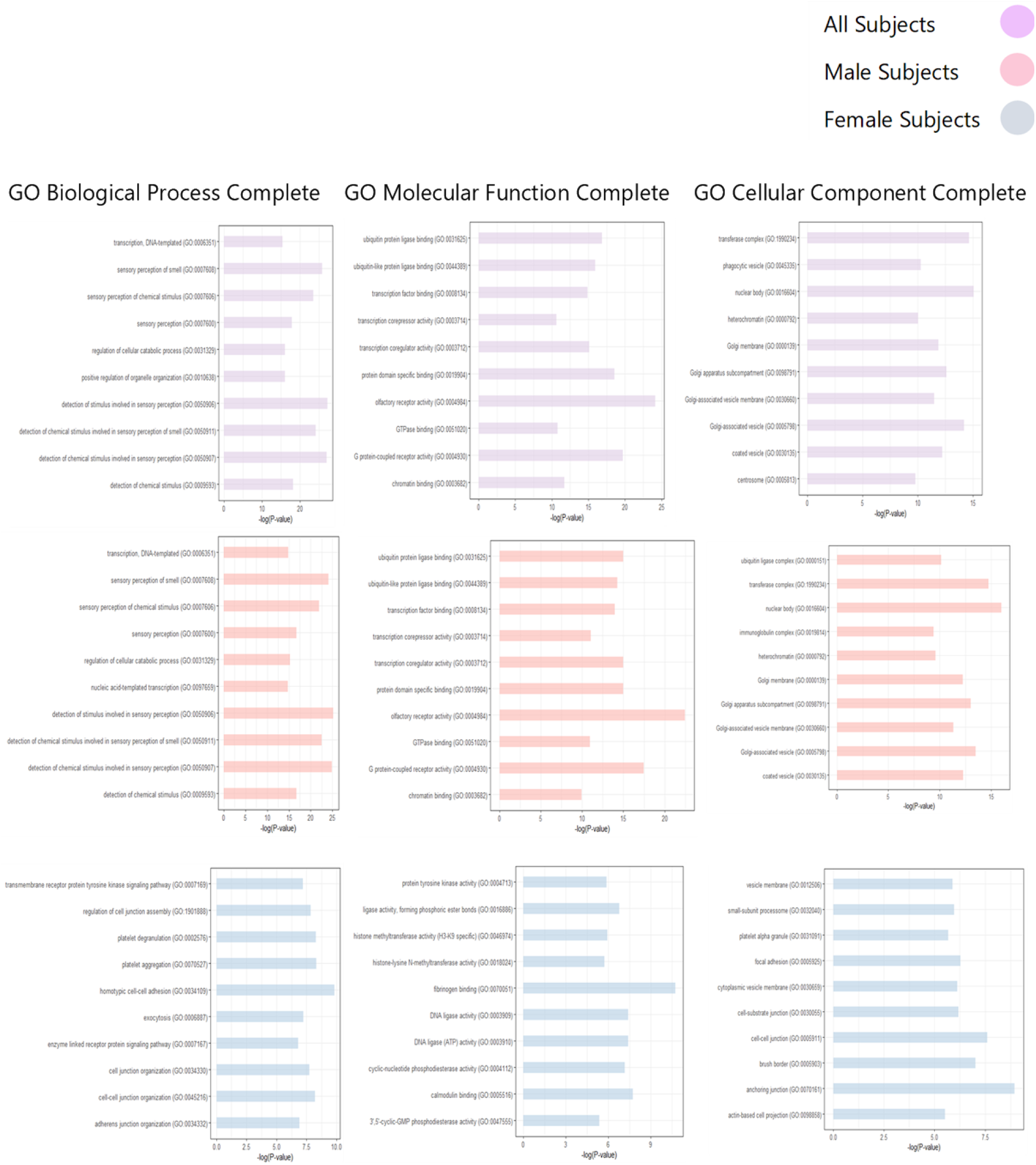
Most Significant Overrepresented PANTHER GO Biological Process, GO Molecular Function, GO Cellular Component Annotations for each pool of personally perturbed genes. The PANTHER GO Ontology database (DOI: 10.5281/zenodo.4081749 Released 2020-10-09) was used to obtain these annotations^17^.

The GO annotations in **Figure 2,** reveal that across all three GO categories, for genes pooled from all patients is nearly identical to the annotations for the genes pooled from all male patients. It is worth noting the cellular locations across all patients, male patients and female patients reveal an overrepresentation of genes localized to vesicle membranes. Across molecular functions transcriptional regulation/repression appears in all patients and male patients. In female patients there is an annotation for histone methyltransferase activity. Modification of histones is a known mechanism by which transcription is regulated^18^. Ubiquitin ligase activity, a top annotation for both all patients and male patients, is interesting as histone ubiquitination is another common histone modification^19^.

The molecular function annotations for both the pool of all and male patient genes have genes overrepresented for GTPase binding and G-protein coupled receptor (GPCR) activity. These annotations are of interest because signaling of neurotransmitters is controlled by GPCR activity^20^. The molecular function annotations for the pool of female patients show genes overrepresented for phosphodiesterase activity. Phosphodiesterases are known to break down cAMP and cGMP^21^. Intracellular signaling in motor circuits, associative/cognitive circuits and limbic circuits have been shown to be mediated by cAMP and cGMP, and as such phosphodiesterase inhibition has become an attractive drug target for neurological disorders^22^. The presence of neurologically relevant annotations for these pools of perturbed genes suggests that the resultant genes are of some significance. The appearance of unnamed genes in these pools of perturbed genes prompted further investigation into their nature and why they might be relevant to ASD etiology.

### Unnamed Genes Appear in Each Study’s Pool of Personally Perturbed Genes

The gene pools for all, male and female patients contain unnamed genes. The same unnamed genes were pulled from the gene pool with no gender filter and the male gender filter; C18orf25, C15orf39, C1orf109, C1orf43, C19orf12, C6orf106, C3orf58, C19orf53, C17orf80, C4orf33, C21 orf2, C10orf2, and C1orf162. The unnamed genes pulled from the female gene pool were C10orf25 and C10orf90. These genes were of interest because of their prevalence in each patient personalized perturbation profile. The prevalence of these 15 unnamed genes in each dataset ranged from 17.1 % to as high as 85.7% of patients **Figure 3**. PubMed results for the male unnamed genes C18orf25, C15orf39, C1orf109, C1orf43, C19orf12, C6orf106, C3orf58, C19orf53, C17orf80, C4orf33, C21orf2, C10orf2, and C1orf162 were relatively scarce with no result count over 200 and ten of the result counts were below 20. The female unnamed genes C10orf25 and C10orf90 were also rarely mentioned in PubMed with result counts below 20 as well.

**Figure 3.**
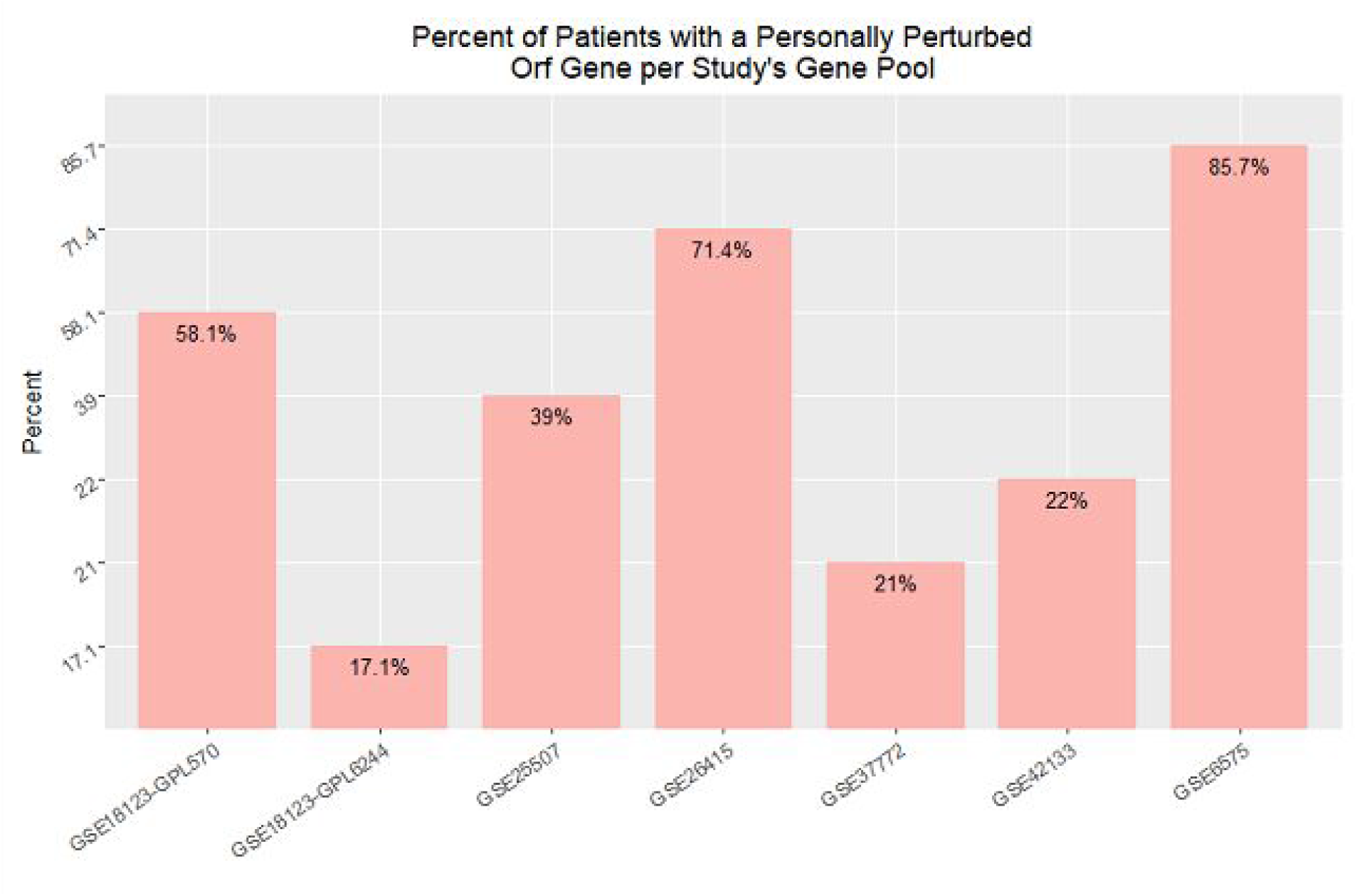
C18orf25, C15orf39, C1orf109, C1orf43, C 19orf12, C6orf106, C3orf58. Ct9orf53, C17orf80, C4orf33, C21orf2, C10orf2, C1orf162, C10orf25 and C10orf90 are personally perturbed in a sizeable fraction of the individuals in each study.

### Differential Correlation Analysis Results

Differential Gene Correlation Analysis was used to identify differentially correlated genes in ASD individuals with the unnamed genes. Unnamed genes identified in the male specific gene pool were tested for correlation against male patients in each study and the same was done for females **Figure 4**. In using differential correlation analysis, we admittingly weaken the strength of any derived correlation, since the datasets used are not only split by disease status but also by gender. It is also true small sample sizes can lead to spurious correlations^23^. However, it should also be mentioned by not separating data, spurious conclusions can also be made^23^. For instance, Aggarwal et al. 2016 notes if hemoglobin levels are plotted against height and not separated by gender, a false correlation will appear^23^. Given the strong male bias in these data and gender-based differences in ASD presentation we thought it pertinent to query correlations after splitting by gender. By dividing by disease status, we get a unique view as to what correlations change between ASD and typically developing populations. Correlations were filtered by the ASD Spearman Correlation Coefficient greater than 0.7 or less than −0.7 and a p-value difference less than 0.05 **Figure 4**. These correlations were filtered by their presence in TCGA Pancancer Low Grade Glioma data where the Spearman Correlation Coefficient was greater than 0.3 and less than −0.3 while also having a p-value less than 0.05 **Figure 4**. The filtered correlations were stored in **Supplemental Data 12**. Using the TCGA Pancancer Low Grade Glioma dataset as a filter was used to safeguard against spurious correlations and the ASD spearman correlation coefficient filter of 0.7 was used to ensure any correlations found were indeed strong. We use TCGA Pancancer Low Grade Glioma data not only for its size and clean data structure, but also because there is evidence of significant abnormalities in the microglia of ASD individuals leading to brain inflammation^24^. Investigation into these correlated genes was aided by the PubMed search tool Chilibot, wherein a pairwise search of gene names was used to derive connections^25^.

**Figure 4.**
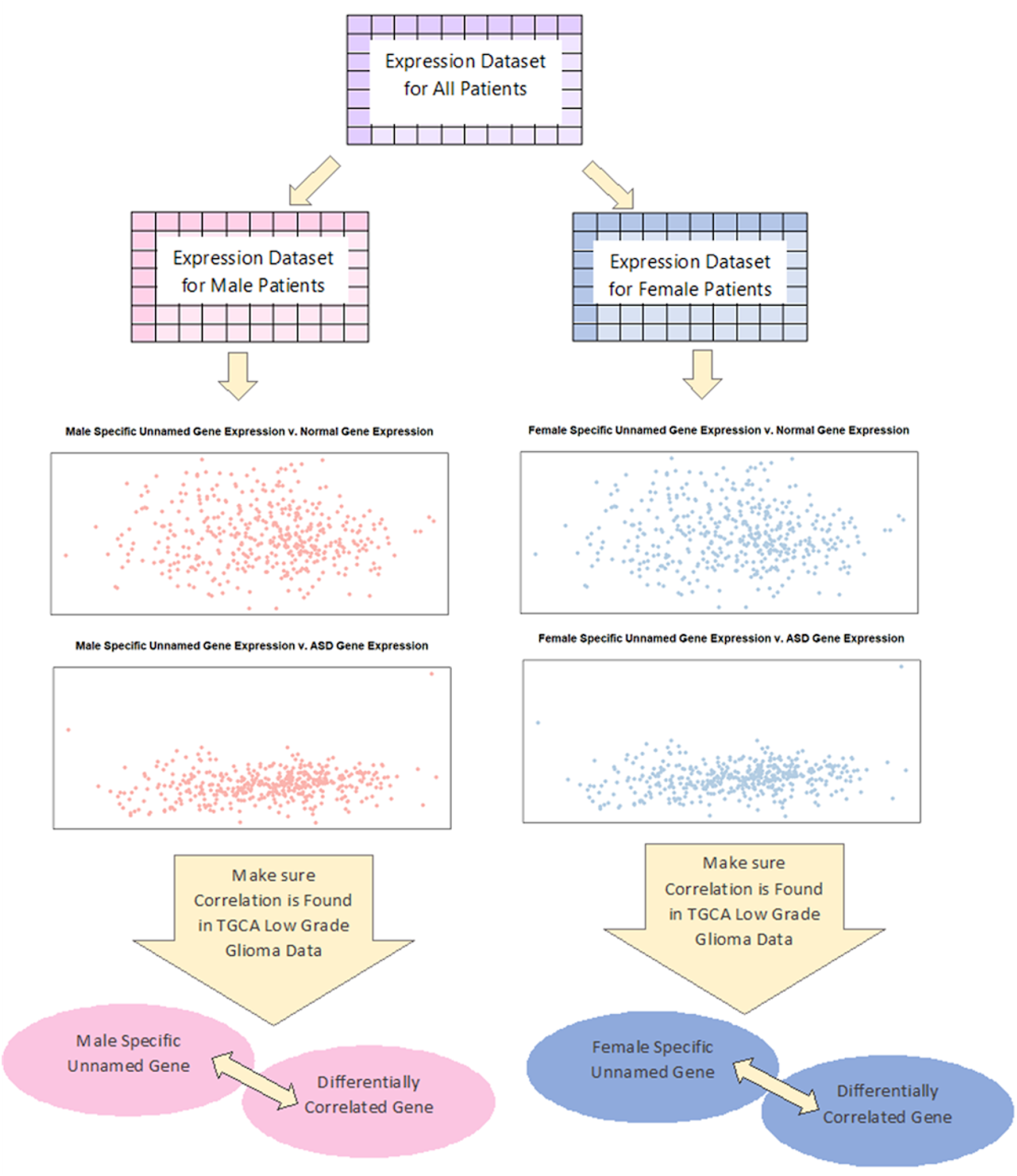
Expression Datasets were separated by gender and Differential Correlation Analysis was performed for male specific unnamed genes and female specific unnamed genes. Correlations were filtered by an ASD Spearman Correlation Coefficient greater than 0.7 and less than −0.7 while also having a p-value difference less than 0.05. These correlations were filtered by their presence in TGCA Pancancer Low Grade Glioma data where the Spearman Correlation Coefficient was greater than 0.3 and less than −0.3 while also having a p-value less than 0.05.

C18orf25, C1orf109, C6orf106, and C19orf53 have been found to localize to the nucleus^26–29^. Two of these genes, C18orf25 and C6orf106 along with C10orf90, have motifs/domains related to ubiquitin. C18orf25 has two small ubiquitin-like modifiers or SUMO motifs and C6orf106 has a ubiquitin-associated UBA-like domain^26,30^ Through the correlation analysis scheme described above we found C18orf25, C1orf109, C19orf12, C6orf106, C3orf58, C19orf53, C4orf33, C10orf2 and C10orf90 are all correlated with a variety of ubiquitin specific peptidases (USPs). USPs are the largest family of deubiquitinating enzymes, noted for their substrate specificity and association with the DNA repair pathway^31^. C10orf90 functions as an E2 ubiquitin ligase involved in responding to DNA damage^32^. C1orf109 is also implicated in DNA damage as it has been shown to lead to the accumulation of an RNA/DNA hybrid structure, called an R-loop, which has been shown to lead to DNA damage^33,34^. The helicase DHX9 was shown to also promote R-loop formation and through the correlation analysis scheme in this work we show it correlates with C10orf90, C17orf80 and C19orf53^35^. It has also been demonstrated that C21orf2 is needed to interact with NEK1 to mitigate DNA damage repair^36^. While not directly associated with DNA damage, C17orf80 is highly correlated with CUL4, a noted E3 ubiquitin ligase, has been found to establish a DNA repair threshold^37^.

C1orf43, C3orf58, and C1orf162 show localization to the Golgi apparatus while^38–40^ C1orf43, C19orf12, C17orf80, C21orf2, C10orf2, have evidence of localization to the mitochondria^38,41–44^. Several of these unnamed genes have ties to phagocytosis/autophagy. By studying L. pneumophila pathogenesis, Jeng et al. 2019 stated C1orf43 as a key regulator of phagocytosis^38^. Venco et al. 2015 propose C19orf12 to be a sensor of mitochondrial damage and inducing autophagy as a protective mechanism to avoid apoptosis^45^. The variant rs3800461 on the gene C6orf1 06 was found by Law et al. 2017 to be implicated in autophagy^46^. Binding of C3orf58/DIPK2A to VAMP7B was shown to encourage autophagosome-lysosome fusion^47^. It is worth noting C3orf58/DIPK2A is directly implicated in familial autism^48^. Biopsies from individuals with mutations in C10orf2 revealed autophagic vacuoles and abnormal mitochondria upon histopathological examination^49^. From the analysis performed in this work additional evidence is provided to support the connection between these poorly characterized genes and autophagy. C18orf25 is highly correlated with JAK2, HIF1A, PIK3C3, RAB10 and BMPR2 all of which are key regulators of autophagy^50–52^. C19orf53 correlates with PARK7, SOD1 and ROCK1 which have roles in autophagic proteolysis and the formation of the autophagosome^53–55^. Our correlation analysis also reveals C1orf109 is highly correlated with TMEM41B, ATG4C RAB39B, TRIM8, and NPRL3 which are also regulators of autophagy^56–59^. C1 orf1 09 was shown here to be highly correlated with SNCA. SNCA, or alpha-synuclein, is implicated in several neurological conditions including dementia and Parkinson’s disease^60^. Minakaki et al. 2018 provided evidence demonstrating components of the autophagy-lysosome pathway are present in extracellular vesicles, extracellular vesicles in cerebrospinal fluid transfer SNCA from cell to cell, and inhibiting the autophagy-lysosomal pathway increases SNCA in neuronal extracellular vesicles^61^.

As mentioned earlier, C19orf12 and C10orf2 have been implicated in mitochondrial function/ disorders^45,49^. Mutations in C19orf12 have made this protein unable to localize to the mitochondrial membrane, causing high levels of mitochondrial Ca^2+^, and an inability to respond to oxidative stress^45^. Mutations in C19orf12 are also shown to be involved with neurodegeneration with brain iron accumulation, a molecular disorder shown to be involved in the disease etiology of frontotemporal dementia, Parkinson’s disease, Alzheimer’s disease, Friedrich ataxia and Amyotrophic lateral sclerosis (ALS)^62^. C10orf2/TWNK is the only mitochondrial helicase needed for mitochondrial DNA replication and its dysfunction is implicated in neurodegeneration and premature aging^63^. As with phagocytosis our correlation analysis scheme has revealed interesting correlations between other unnamed genes and the mitochondria. C1orf43 correlates with VDAC1, TOMM20, and TMCO1 whose dysfunction/overexpression has been linked to mitophagy and mitochondrial dysfunction in relation to Alzheimer’s disease and Parkinson’s Disease^64–66^. It is worth mentioning the two unnamed genes with direct connections to mitochondrial dysfunction, C19orf12 and C10orf2, are correlated with TMCO1 and TOMM20, respectively.

## Discussion

To the best of our knowledge this is the first-time personalized perturbation profiles, as laid out by Menche et al. 2017, have been used to establish concordance among blood based ASD transcriptomic datasets. Using the Welch two sample t-test we saw only one gene, SMARCA2, was differentially expressed between all datasets. By leveraging personalized perturbation profiles, we find each study’s pool of perturbed genes, male perturbed genes and female perturbed genes have 1,189, 1,111, and 133 genes in common, respectively. PANTHER GO annotations of these gene pools reveal interesting connections to vesicle membranes, GPCRs, and phosphodiesterase activity. Extracellular vesicles have recently been implicated in the disease etiology of ASD while GPCRs/phosphodiesterases are an important facet of neural signaling^20,21,67^.

In these pools of perturbed genes, we found 15 unnamed genes which had a relatively high prevalence among each study’s pool of perturbed genes. Of these, 13 unnamed genes were common to pools of each study’s male pool of perturbed genes and two of these were common to each study’s female pool of perturbed genes – the discrepancy likely owing to the smaller number of female patients. Investigating these poorly characterized genes along with differential correlation analysis demonstrated overlap between their correlations and experimentally derived functions. We find these unnamed genes have connections with DNA damage, ubiquitination, autophagy, and the mitochondria.

Mitochondrial dysfunction has been implicated in ASD. Magnetic Resonance Spectroscopy has shown N-acetyl-aspartate, a mitochondrial dysfunction marker, was significantly lower in ASD children^68^. Aberrations in respiratory chain complexes have also been found in ASD individuals^69^. Similarly, elevated levels of reactive oxygen species were observed in the brains of ASD subjects^70^. Extracellular vesicles in ASD microglia were shown to have significantly more mitochondrial DNA than typically developing individuals^67^. A recent study by Varga et al. 2018 found over 16% of ASD individuals tested in their study had deletions of mitochondrial DNA^71^. Mitochondrial mutations may also provide an answer as to why ASD seems to affect more males than females. Frank 2012 presented evidence that mitochondrial mutations act as a sex-biased sieve allowing males to build up more mitochondrial mutations^72^. Support for this theory is provided by a study using *Drosophila Melanogaster,* in which mitochondrial mutations disproportionally affect male aging as opposed to female aging^73^. Siddiqui et al. 2016 acknowledges while mitochondrial dysfunction is not yet designated a causal factor for ASD, indirect evidence for this hypothesis is rising^69^.

Oxidative stress has also been related to DNA damage in relation to neurological disorders. Melnyk et al. 2012 found evidence of depleted antioxidant levels and oxidative DNA damage in the plasma of ASD individuals^74^. Markkanen et al. 2016 remarked that studies are starting to implicate DNA repair mechanisms in ASD but at times have provided conflicting results^75^. In our results we noticed C1orf109 is implicated in DNA damage through the accumulation of R-loops and C10orf90, C17orf80 and C19orf53 were highly correlated with DHX9, a protein known to promote R-loop formation^35^. In the context of neurological diseases, embryonic neural R-loop levels were determined to be essential for proper nervous system development^76^. Defective R-loop mechanisms have also been implicated in the molecular workings of ALS^77^. What is more, is evidence provided by Akman et al. 2016 show mutations in RNAase H1 have proved detrimental to mitochondrial R-loop levels, which have led to mitochondrial DNA aggregates^78^. Holt 2019 remarks this atypical organization of mitochondrial DNA is implicated in infantile-onset epilepsy, mitochondrial encephalopathy, and cerebellar dysfunction^79^. This finding might prove a useful connection between R-loops, DNA damage, mitochondrial dysfunction, and the presence of mitochondrial DNA in extracellular vesicles.

Autophagy is also starting to have a more defined role in the molecular mechanics of ASD. Microglia with defunct autophagic pathways have been shown to negatively affect synaptic pruning in ASD individuals^80^. Recently, Dana et al. 2020 found the autophagic markers LC3 and Beclin-1 were significantly disturbed in the Cc2d1a animal model of ASD^81^. What is interesting about this study is their separation by gender revealed while both ASD males and females had lower levels of Beclin-1, LC3 expression was increased in ASD females and decreased in ASD males^81^. This finding seems indicative of gender specific autophagic responses. Kasherman et al. 2020 detailed deficits in the mTOR signaling, a method by which autophagy is regulated, is disturbed in ASD subjects, and collected evidence also linking mutations in USPs to ASD features^82–84^. Kasherman et al. 2020 also states the role of USPs in ASD etiology remains poorly studied^82^. A possible answer might be found in Klusmann et al. 2018, as they provided an interesting connection between ubiquitination and R-loops, being ubiquitination of histones aided in the suppression of R-loops^85^.

More experimental validation must be done to provide conclusive links between DNA damage, R-loops, autophagy, mitochondrial damage, and ubiquitination in the context of ASD; our research here only draws attention to a handful of genes that may have been overlooked as they are poorly studied, and in the case of R-Loops, represent a relatively new field of study. The inclusion of more females in ASD blood transcriptomics would be of enormous value as well. As stated earlier, these data are derived from over 90% male ASD subjects and as such our analysis of the ASD female blood transcriptome is limited. However, we find that useful information can still be gleaned from these data with the use of personalized perturbation profiles. Aside from collecting a common pool of perturbed genes, which the t-test could not deliver, we found poorly characterized genes in those pools which may underlie ASD molecular mechanics. These results are a promising step in cataloging the true heterogeneity of ASD.

## Materials and Methods

### ASD Blood-Based GEO Datasets

To obtain relevant datasets we queried GEO with the terms “ASD” or “autism”. These results were limited to human samples and only those of blood-based tissue types^86^. We filtered these results further by only examining studies with more than 20 ASD individuals. Expression, patient, and probe data for each study was obtained via the R package GEOquery^87^. If the expression data was not already normalized the data was normalized by adding the minimum value of the expression dataset, taking the log2 of these data and normalizing by quantiles by use of the R package limma^88^.

### Personalized Perturbation Profiles and T-test

Each expression dataset was split by disease status and subjects who were not labeled as ASD were excluded from analysis. The t-test p-value was calculated using the Welch two sample t-test for each gene, with the expression of the control group being compared against the expression of the ASD group. Genes with a p-value below 0.05 were added to each study’s list of differentially expressed genes. All these studies were compared with one another to obtain common differentially expressed genes. Establishing personalized perturbation profiles began with obtaining the mean and standard deviation of the expression of each gene in the control group. The z-score, for each ASD individual at each gene was obtained by the following equation as laid out by Menche et al. 2017^13^:

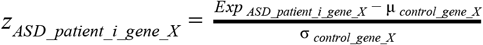

Where, *z_ASD_patient_t_gene_x_* is the z-score for ASD patient i at gene X, *Exp_ASD_patient_i_gene_X_* is the expression of ASD patient i at gene X μ_*control_gene_X*_ is the mean expression for the control group at gene X, and σ_*control_gene X*_ is the standard deviation of the expression of the control group at gene X. A gene was considered perturbed if the z-score was greater than 2.5 or less than −2.5. In this fashion each patient had a list of genes which were determined personally perturbed. These genes were added to each study’s pool of personally perturbed genes. These pools of perturbed genes were compared with one another to obtain a pool of commonly perturbed genes. The common male personally perturbed gene pool was obtained by limiting each study’s pool of personally perturbed genes to male patients, and these male only study pools were compared with one another. Common female specific personally perturbed genes were obtained in the same way.

### PANTHER Enrichment

Pools of personally perturbed genes were run through the PANTHER Overrepresentation Test GO Biological Process Complete, GO Molecular Function Complete, and GO Cellular Component Complete utilizing the GO Ontology database DOI: 10.5281/zenodo.4081749 Released 2020-10-09^17^. The resulting data tables were downloaded and were filtered to exclude any annotation with more than 1000 genes linked to it. The negative log was taken of the raw p-value and only the top 10 most significant annotations were displayed. Plots were generated using the R package ggplot2 and colors were obtained using the R package RColorBrewer^89,90^. These packages were also used when plotting the prevalence of unnamed genes in each study’s pool of perturbed genes.

### Differential Correlation Analysis and Chilibot

Differential correlation analysis was performed using the R package DGCA^91^. Design matrices were created to split data based on ASD status. The spearman correlation coefficient used as a measure of differential correlation. Differential correlation was only obtained for the unnamed genes found in the pools of commonly perturbed genes. Only those genes with a p-value difference less than 0.05 and an ASD spearman correlation coefficient greater than 0.7 and less than −0.7 were explored further. As a level of noise reduction TCGA Pancancer Low Grade Glioma Coexpression data was used to verify correlations found by differential correlation analysis. Coexpression data mRNA Expression, RSEM Batch normalized from Illumina HiSeq_RNASeqV2, for the unnamed genes was downloaded from cBioPortal^29–92–101^. These coexpression data were filtered to exclude correlations whose p-value is above 0.05 and whose spearman correlation coefficient is below 0.3 and above −0.3 but still below 0. Those correlations not found in these refined TCGA Pancancer Low Grade Glioma data were excluded from further analysis. The resulting correlated genes were investigated using the PubMed text mining tool, Chilibot^25^. Chilibot’s pairwise search function was used to determine how correlated genes might be connected.

## Supplementary Data

Supplementary Data are available at BioRxiv online.

